# Photoreceptor Compartment-Specific TULP1 Interactomes

**DOI:** 10.1101/2021.06.07.447411

**Authors:** Lindsey A. Ebke, Satyabrata Sinha, Gayle J.T. Pauer, Stephanie A. Hagstrom

## Abstract

Photoreceptors are highly compartmentalized cells with large amounts of proteins synthesized in the inner segment (IS) and transported to the outer segment (OS) and synaptic terminal. Tulp1 is a photoreceptor-specific protein localized to the IS and synapse. In the absence of Tulp1, several OS-specific proteins are mislocalized and synaptic vesicle recycling is impaired. To better understand the involvement of Tulp1 in protein trafficking, our approach was to physically isolate Tulp1-containing photoreceptor compartments by serial tangential sectioning of retinas and to identify compartment-specific Tulp1 binding partners by immunoprecipitation followed by liquid chromatography tandem mass spectrometry. Our results indicate that Tulp1 has two distinct interactomes. We report the identification of: 1) an IS-specific interaction between Tulp1 and the motor protein Kinesin family member 3a (Kif3a), 2) a synaptic-specific interaction between Tulp1 and the scaffold protein Ribeye, and 3) an interaction between Tulp1 and the cytoskeletal protein Microtubule-associated protein 1B (MAP1B) in both compartments. Immunolocalization studies in the wild-type retina indicate that Tulp1 and its binding partners co-localize to their respective compartments. Our observations are compatible with Tulp1 functioning in protein trafficking in multiple photoreceptor compartments, likely as an adapter molecule linking vesicles to molecular motors and the cytoskeletal scaffold.

## 1. Introduction

Retinal photoreceptor cells are an extraordinary example of cells containing morphologically distinct compartments, coordinating proteins to optimize each specialized function [1]. The outer segment (OS) compartment is considered a modified primary cilium packed with a series of discrete disc membranes which contain the photopigment molecules and accessory proteins necessary for the phototransduction cascade [2,3]. The OSs are isolated from the rest of the cell by a narrowed counterpart to the transition zone, called the connecting cilium (CC), which is a site of selective protein transport. The inner segment (IS) compartment houses all of the biosynthetic machinery required for protein synthesis, as well as the mitochondrial powerhouse of the cell. Distal to the ISs are rows of photoreceptor cell bodies containing the nucleus followed by the outer plexiform layer (OPL) where synaptic termini make contact with second order neurons through a specialized ribbon structure, eventually transmitting signals centrally to the brain for processing into a visual image.

The development, maintenance and function of photoreceptor sensory cilia require the transport of both cilium-specific and OS-specific proteins from the IS to the appropriate cellular compartment. This protein transport occurs throughout the life of the photoreceptor as the OSs are continuously renewed approximately every ten days [4,5]. This rapid OS turnover requires an extraordinary high rate of protein synthesis and efficient protein trafficking. Photoreceptors use several different mechanisms to isolate their cellular compartments from one another and to guide the delivery of proteins via vesicular transport [1]. Intraflagellar transport (IFT) is recognized as an essential process in photoreceptors by which proteins are bidirectionally transported along microtubules. The IFT protein complex is arranged into two particles functioning as adaptors between molecular motors and cargo [6]. This movement is facilitated by two different motor complexes - the heterotrimeric Kinesin-II complex which transports the IFT-B complex anterograde or toward the plus-end of the microtubule and the cytoplasmic Dynein-2 complex which transports the IFT-A complex retrograde or toward the minus-end of the microtubules. Rhodopsin constitutes the majority of the OS-resident protein and therefore vectorial transport of this protein has been studied in great detail. Studies indicate distinct mechanisms of rhodopsin trafficking, including IFT-dependent movement, IFT-independent movement and regulation via Arf and Rab GTPases [7,8]. Photoreceptors also use several other protein trafficking mechanisms to coordinate functional compartmentalization including distinct plasma membrane composition, adaptor proteins, chaperones and targeting signals encoded within protein sequences [1]. It is clear that there are several unique processes driving intrinsic protein compartmentalization; however, critical knowledge gaps still remain regarding the specific pathways and components involved. Even less is known about protein trafficking to the synaptic terminal and assembly of the specialized ribbon structure associated with large numbers of synaptic vesicles primed for neurotransmission. A few studies have shown that presynaptic proteins are transported as “packages” to synaptic sites consisting of aggregates of separate membranous organelles and proteins [9,10]. The identities of the molecular machinery and the precise regulatory mechanisms underlying trafficking of synaptic components remain unclear.

Tubby like protein 1 (Tulp1) is a photoreceptor-specific protein localized to the IS, CC and synaptic termini and linked to protein trafficking [11–13]. Phenotypic evidence from *tulp1-/-* mice indicates an early-onset and rapid photoreceptor degeneration with loss of retinal function [11–13]. A detailed analysis of *tulp1-/-* photoreceptors revealed that rhodopsin, cone opsins and several other OS phototransduction proteins are mislocalized and distributed throughout all photoreceptor compartments, prior to degeneration [11,14]. Interestingly, the differential distribution of OS-resident proteins implies a role for Tulp1 in the transport of specific proteins but not others, supporting the notion of multiple ciliary transport routes [15–17]. Since Tulp1 is also localized to photoreceptor synaptic terminals, further investigation of *tulp1-/-* retinas revealed that the synapses lack the tight spatial relationship between the ribbon-associated proteins, Bassoon and Piccolo, indicating a disruption in the continuous vesicular cycling of neurotransmitter release [14].

Identifying the proteins involved in regulating and maintaining compartmentalization and accurate protein transport is vital to understanding the function of photoreceptor cells. The unique phenotype of the *tulp1-/-* retina characterized by distinct defects in multiple photoreceptor compartments suggests that Tulp1 may play different roles at opposite ends of the cell. To better understand the involvement of Tulp1 in protein trafficking, our approach in this study was to physically separate and isolate the Tulp1-containing photoreceptor compartments and characterize unique binding partners. Herein we report the identification of: 1) an IS-specific interaction between Tulp1 and the motor protein Kinesin family member 3a (Kif3a), 2) a synaptic-specific interaction between Tulp1 and the scaffold protein Ribeye, and 3) an interaction between Tulp1 and the cytoskeletal protein Microtubule-associated protein 1B (MAP1B) in both compartments.

## 2. Results

### 2.1. Photoreceptor compartment isolation

We took advantage of the layered structure of the retina where distinct subcellular compartments are well-defined and isolated the Tulp1-containing compartments by serial tangential sectioning of the retina. Prior to further analysis, we evaluated the quality of the retina and the quality of the serial sectioning using three methods. First, each retinal flat mount preparation was imaged by optical coherence tomography (OCT). The OCT provides information about how intact the retinal layers remain after dissection and manipulation. Figure 1A shows the B-scan view of a representative preparation revealing a flat retina with a slight defect on the far left side, highlighted by a star. Any disturbed or folded peripheral edges were removed prior to tangential sectioning. The second method used to assess the quality of the tangential sectioning was a dot blot of all serial sections probed with antibodies against rhodopsin. A blot in which only the outermost 4-5 sections corresponding to the photoreceptor OSs were strongly positive for rhodopsin indicated optimally sectioned retinas (Figure 1B). And lastly, the protein content of each tangential section was analyzed by Western blot. Figure 1C shows a representative Western blot of an experimental “set”, defined as serial sections 1 – 27, probed for known retinal compartment-specific proteins.

**Figure 1.**
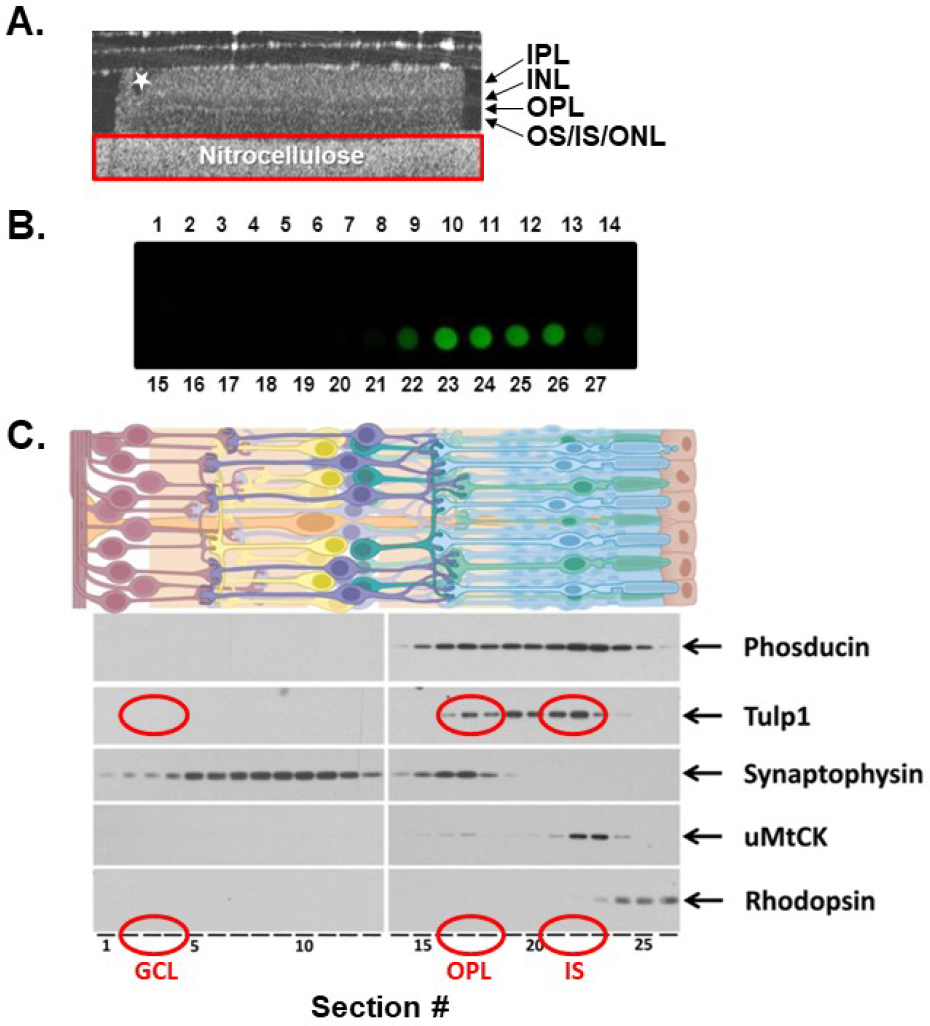
Isolation of Tulp1-containing photoreceptor compartments. (A) OCT image of a retinal flat mount preparation to assess the quality of the dissected retina. Any disturbed or folded peripheral edges, highlighted by a star, were removed prior to tangential sectioning. The red rectangle encompasses the nitrocellulose support. (B) A dot blot analysis of all serial sections probed with antibodies against rhodopsin. A blot in which only the outermost 4-5 sections corresponding to the photoreceptor OSs were strongly positive for rhodopsin indicated optimally sectioned retinas. (C) A schematic illustration of retina showing each cell type. The protein content of each tangential section was analyzed by Western blot probed for known compartment-specific proteins. The five marker proteins included Phosducin, Tulp1, Synaptophysin, uMtCK and Rhodopsin. Each lane of the gel represents the protein content of a single 10 μm section starting from the GCL and progressing through the photoreceptor OSs. The GCL-, IS-, OPL-containing samples used for further analysis are circled in red.

The following proteins were used to verify the spatial resolution of each compartment. Phosducin is a photoreceptor-specific soluble protein known to interact with the βγ subunits of G proteins and is evenly distributed throughout the entire photoreceptor cytoplasm [18]. Tulp1 is restricted to the photoreceptor CC, IS and synapse [11]. Synaptophysin is a marker for pre-synaptic vesicles expressed in the photoreceptor terminals of the OPL and also in the second order neuron terminals of the inner plexiform layer (IPL) [19]. Ubiquitous mitochondrial creatine kinase is an enzyme critical to mitochondrial respiration and is localized in the photoreceptor IS and synapse [20]. Finally, rod OSs were identified by the presence of rhodopsin. Results indicate that our experimental design provides an accurate isolation of photoreceptor compartment-specific regions and protein distribution within the retina (Figure 1).

Following serial sectioning and compartment isolation, three sections from the IS compartment were combined and three sections from the OPL were combined to provide a sufficient amount of protein for further analysis. As negative controls, an equal number of serial sections from a retinal region lacking Tulp1, such as the ganglion cell layer (GCL), as well as an equal number of serial sections from *tulp1-/-* photoreceptors were collected.

### 2.2. Tulp1 photoreceptor compartment-specific binding partners

To identify Tulp1 interacting proteins, we performed immunoprecipitations (IPs) from the photoreceptor compartment-specific lysates using anti-Tulp1 antibodies. Compartment-specific IP products were analyzed by liquid chromatography tandem mass spectrometry (LC MS/MS). We used an Orbitrap system due to its increased sensitivity, speed, and ability to resolve peptides from a complex mixture [21]. Three separate IPs were performed and combined proteomic results revealed 110 potential Tulp1 binding partners unique to the IS, 17 proteins unique to the synapse, and 177 proteins common to both compartments. To better analyze this data set, we categorized the proteins into functional groups using the Universal Protein Resource (UniProt) online database. Categories containing the most proteins from the IS analysis and the combined compartment analysis (IS and OPL) included heat shock proteins, ribosomal proteins, and histones. This is not surprising given that the IS is the site of protein synthesis; however, the majority of these likely represent non-specific interactions as they are members of the most frequently detected protein families in proteomic experiments [22]. This was verified employing a web-based resource called the contaminant repository for affinity purification mass spectrometry data, the “CRAPome”, which stores and annotates negative control data generated by the proteomics community (http://www.crapome.org).

To further refine our data, stringent criteria were applied in order for proteins to be considered potential Tulp1 binding partners: 1) proteins must have five or more spectral counts, 2) proteins must be identified by at least two unique peptides, 3) proteins must have a greater than three-fold increase in spectral counts compared to negative controls, and 4) proteins must be identified in at least two of the three independent IP experiments. As an internal control, Tulp1 was identified in the IS compartment by 48 unique peptides, corresponding to 63% sequence coverage; and Tulp1 was identified in the synaptic compartment by 28 unique peptides, corresponding to 38% sequence coverage. Following these criteria, there were a limited number of proteins in each compartment analysis that exhibited the strongest proteomic results. We chose to validate these observed proteins. In the IS, the candidate proteins include Microtubule-associated protein 1B (MAP1B) and Kinesin family member 3a (Kif3a). In the photoreceptor synapse, the candidates include MAP1B and Ribeye.

### 2.3. MAP1B is a Tulp1-interacting partner in the IS and OPL

Microtubule-associated protein 1B (MAP1B) was identified as a strong candidate Tulp1 interactor in both the IS and OPL compartments. MAP1B is a neuronal-specific microtubule binding protein important for axonal growth and synaptic maturation [23]. In photoreceptors, MAP1B is involved in regulating motor-cargo interactions along the axoneme and in recruiting vesicles to the synaptic ribbon surface [23–28].

We performed a reciprocal IP of rat retinal lysate using antibodies against MAP1B. Figure 2 shows the Western blot analysis of the IP samples probed with Tulp1 antibodies. A band corresponding to the correct molecular weight of Tulp1 (~75 kDa) is detected in the MAP1B IP product lane but not in the IP control lane using non-specific IgG antibodies. As positive and negative controls, Tulp1 is detected in both wt mouse and rat retinal lysate and is not detected in *tulp1-/-* retinal lysate. To verify the efficacy of the IP, the blot was probed for MAP1B. A band corresponding to the correct molecular weight of MAP1B (~320 kDa) is detected in the specific MAP1B IP product, wt mouse and rat retinal lysate, and *tulp1-/-* retinal lysate lanes. Liver lysate was included as a separate negative control for both proteins. The density of the band remaining in the unbound MAP1B IP product lane suggests that only a fraction of MAP1B binds to Tulp1 *in vivo*.

**Figure 2.**
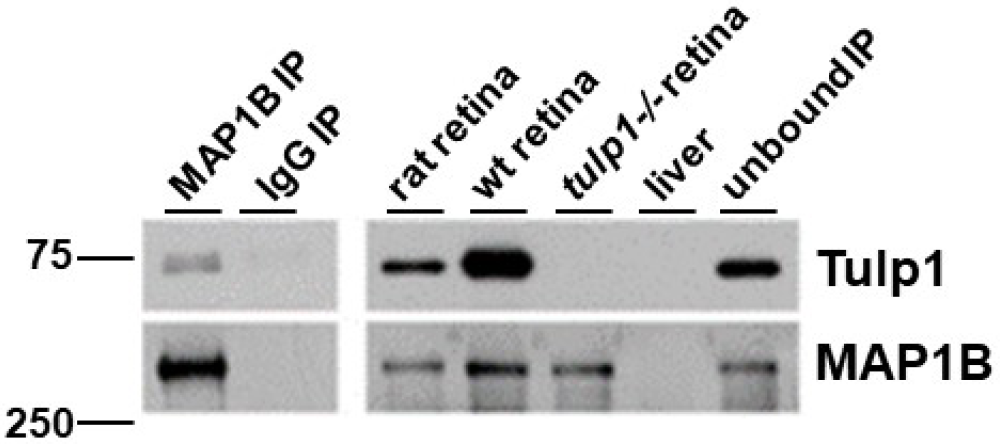
IP of retinal lysate. The top panels show Western blot analysis of the MAP1B IP experimental samples probed with Tulp1 antibodies. In the IP product lane, a band corresponding to Tulp1 is detected. A corresponding band is seen in the rat retinal lysate and wt mouse retinal homogenate but not in the *tulp1-/-* retinal lysate, liver lysate or non-specific IgG IP lanes. The bottom panels show Western blot analysis of the MAP1B IP experimental samples probed with MAP1B antibodies. In the IP product, rat retinal lysate, wt mouse retinal lysate and *tulp1-/-* retinal lysate lanes, a band corresponding to MAP1B is detected. No bands are seen in the liver or non-specific IgG IP sample lanes.

Next, we examined the photoreceptor localization of MAP1B in wt compared to *tulp1-/-* retinal sections by immunohistochemistry (IHC). All localization studies were conducted at P17, an age at which all cell types of the retina are present in wt mice, but which precedes photoreceptor cell death in *tulp1-/-* mice [11,14,15]. In the wt retina, Figure 3A shows MAP1B staining in the OPL and throughout the perikarya and IS of the photoreceptor cell layer. As previously reported, Tulp1 is present in the OPL, perikarya and IS of the photoreceptors [11]. High magnification confirms that both MAP1B and Tulp1 co-localize to the IS and OPL of photoreceptor cells. Figure 3B indicates that the localization of MAP1B in the *tulp1-/-* retina appears similar to that in the wt retina with immunoreactivity seen in the OPL, perikarya and IS. These results imply that the absence of Tulp1 does not grossly affect the photoreceptor distribution of MAP1B.

**Figure 3.**
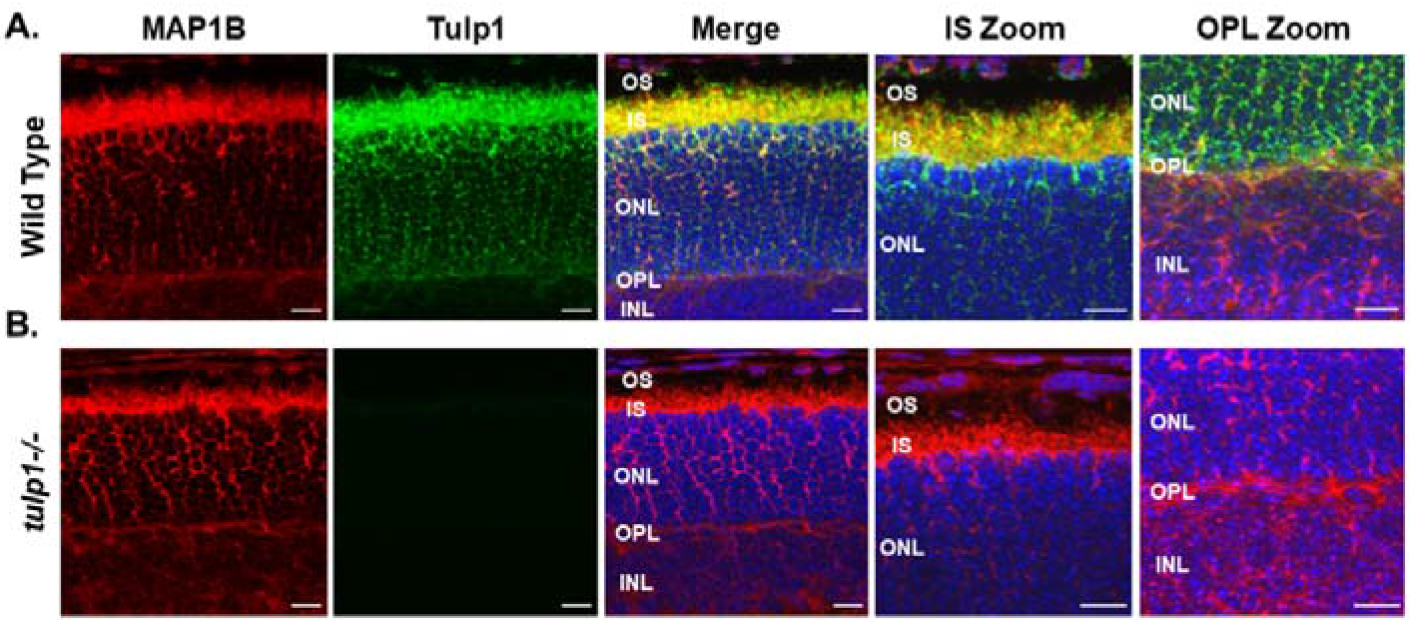
Immunolocalization of MAP1B and Tulp1 in P17 mouse retinas. (A) Wt retinal sections stained with MAP1B (red) and Tulp1 (green). (B) *Tulp1-/-* retinal sections stained with MAP1B (red) and Tulp1 (green). Sections were counterstained with DAPI (blue). Scale bar: 50 μm. INL, inner nuclear layer; OPL, outer plexiform layer; ONL, outer nuclear layer; IS, inner segment layer; OS, outer segment layer.

### 2.4. Kif3a is a Tulp1-interacting partner in the IS

The second protein we confirmed as an IS Tulp1 interacting partner was the Kif3a subunit of Kinesin-2. Heterotrimeric Kinesin-2 is a plus-end directed molecular motor that is required for the anterograde movement of proteins along microtubules within cilia [29]. It consists of two motor subunits, encoded by the Kif3a and Kif3b genes, and an accessory non-motor subunit, encoded by Kap3 [25]. In photoreceptors, in addition to ciliary protein trafficking, Kif3a is critical for maintenance of the axoneme [30,31].

A reciprocal IP of rat retinal lysate using antibodies against Kif3a and probed for Tulp1 detects the correct molecular weight band for Tulp1 in the Kif3a IP lane but not in the IP control lane using non-specific IgG antibodies (Figure 4). As positive and negative controls, Tulp1 is detected in both wt mouse and rat retinal lysate and is not detected in *tulp1-/-* retinal lysate. To verify the efficacy of the IP, the blot was probed for Kif3a. Figure 4 shows that a band corresponding to the correct molecular weight of Kif3a (~80kDa) was detected in the specific Kif3a IP product, rat retinal lysate, mouse wt retinal lysate, and *tulp1-/-* retinal lysate lanes. Heart lysate was included as a negative control for both proteins. The density of the band remaining in the unbound Kif3a IP product lane suggests that only a fraction of Kif3a binds to Tulp1 *in vivo*.

**Figure 4.**
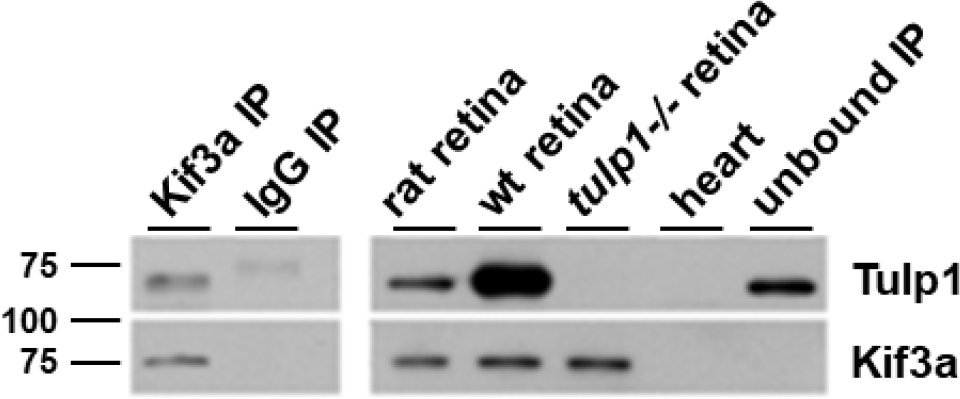
IP of retinal lysate. The top panels show Western blot analysis of the Kif3a IP experimental samples probed with Tulp1 antibodies. In the IP product lane, a band corresponding to Tulp1 is detected. A corresponding band is seen in the rat retinal lysate and wt mouse retinal homogenate but not in the *tulp1-/-* retinal lysate, heart lysate or non-specific IgG IP lanes. The bottom panels show Western blot analysis of the Kif3a IP experimental samples probed with Kif3a antibodies. In the IP product, rat retinal lysate, wt mouse retinal lysate and *tulp1-/-* retinal lysate lanes, a band corresponding to Kif3a is detected. No bands are seen in the heart or non-specific IgG IP sample lanes.

Next, we examined the photoreceptor localization of Kif3a in wt compared to *tulp1-/-* retinal sections by IHC. In wt photoreceptors at P17, Kif3a staining is present in the OPL, perikarya and ISs (Figure 5A). As previously reported, Tulp1 is present in the OPL, perikarya and IS of photoreceptors [11]. High magnification confirms that both Kif3a and Tulp1 co-localize to the IS of photoreceptor cells. Figure 5B indicates that the localization of Kif3a in the *tulp1-/-* retina appears similar to that in the wt retina with immunoreactivity seen in the OPL, perikarya and IS. These results imply that the absence of Tulp1 does not grossly affect the photoreceptor distribution of Kif3a.

**Figure 5.**
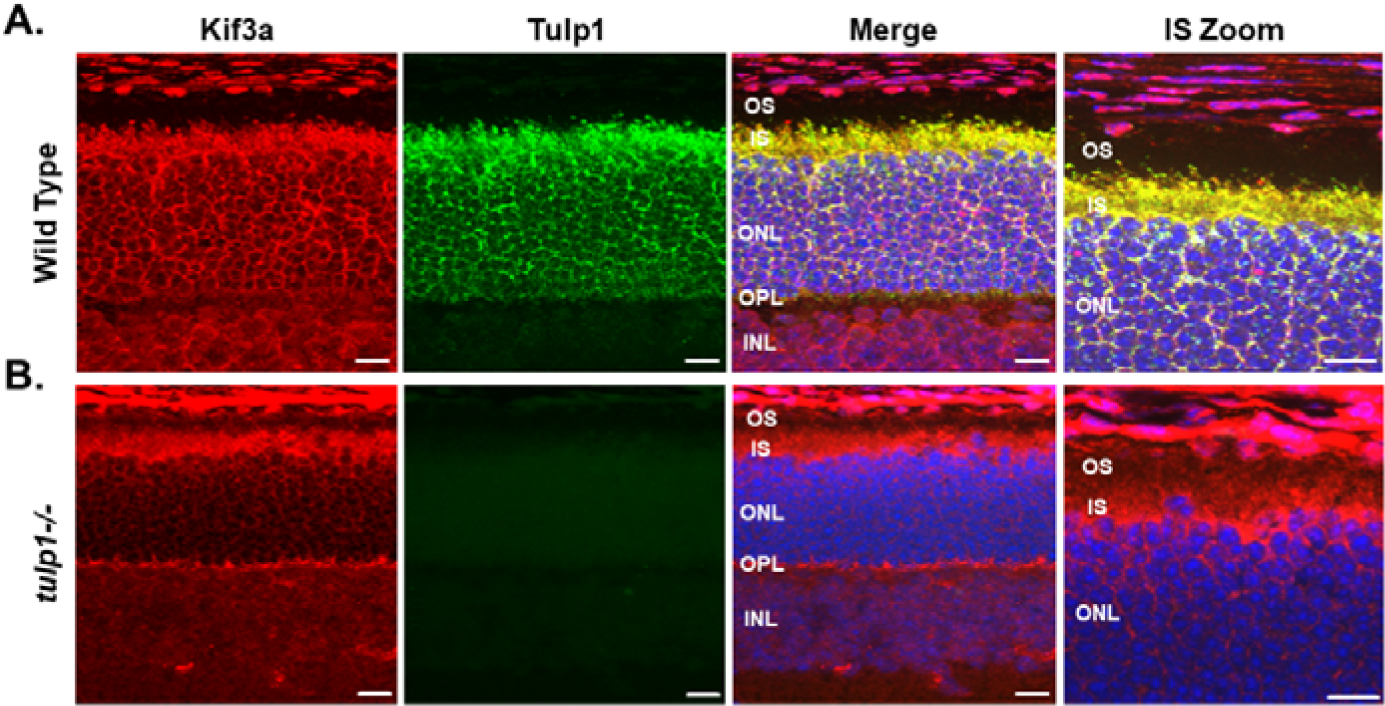
Immunolocalization of Kif3a and Tulp1 in P17 mouse retinas. (A) Wt retinal sections stained with Kif3a (red) and Tulp1 (green). (B) *Tulp1-/-* retinal sections stained with Kif3a (red) and Tulp1 (green). Sections were counterstained with DAPI (blue). Scale bar: 50 μm. INL, inner nuclear layer; OPL, outer plexiform layer; ONL, outer nuclear layer; IS, inner segment layer; OS, outer segment layer.

### 2.5 Ribeye is a Tulp1-interacting partner in the OPL

The second protein we confirmed as an OPL Tulp1 interacting partner was Ribeye, the primary structural protein of photoreceptor synaptic ribbons. In photoreceptors, the synaptic ribbon is a large, plate-like structure that is anchored to the presynaptic membrane and characterized by numerous tethered vesicles that facilitate continuous vesicle release critical for neurotransmission [32]. Ribeye is unique in that it is the only known protein specific to synaptic ribbons [33].

Figure 6 is a reciprocal IP of rat retinal lysate using antibodies against Ribeye that shows Tulp1 in the Ribeye IP lane but not in the IP control lane. As positive and negative controls, Tulp1 is detected in both wt mouse and rat retinal lysate and is not detected in *tulp1-/-* retinal lysate. To verify the efficacy of the IP, the blot was probed for Ribeye. A band corresponding to the correct molecular weight of Ribeye (~120 kDa) is detected in the specific Ribeye IP product, rat retinal lysate, mouse wt retinal lysate and *tulp1-/-* retinal lysate lanes. Spleen lysate was included as a negative control for both proteins. The density of the band remaining in the unbound Ribeye IP product lane suggests that only a fraction of Ribeye binds to Tulp1 *in vivo*.

**Figure 6.**
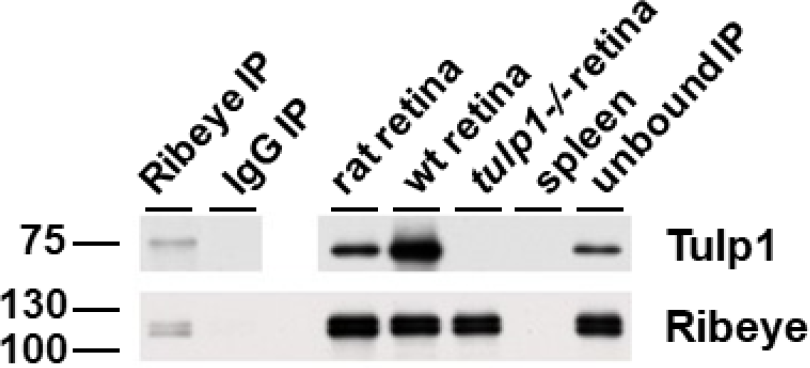
IP of retinal lysate. The top panels show Western blot analysis of the Ribeye IP experimental samples probed with Tulp1 antibodies. In the IP product lane, a band corresponding to Tulp1 is detected. A corresponding band is seen in the rat retinal lysate and wt mouse retinal homogenate but not in the *tulp1-/-* retinal lysate, spleen lysate or non-specific IgG IP lanes. The bottom panel shows Western blot analysis of the Ribeye IP experimental samples probed with Ribeye antibodies. In the IP product, rat retinal lysate, wt mouse retinal lysate and *tulp1-/-* retinal lysate lanes, a band corresponding to Ribeye is detected. No bands are seen in the spleen or non-specific IgG IP sample lanes.

Next, we examined the photoreceptor localization of Ribeye in wt compared to *tulp1-/-* retinal sections by IHC at P17. In the wt retina, Figure 7A shows that Ribeye is localized only to the photoreceptor OPL. As previously reported, Tulp1 is present in the OPL, perikarya and IS of the photoreceptors [11]. High magnification confirms that both Ribeye and Tulp1 co-localize to the OPL. Figure 7B indicates that the localization of Ribeye in the *tulp1-/-* retina is comparable to that in the wt retina with immunoreactivity seen only in the OPL. These results imply that the absence of Tulp1 does not grossly affect the photoreceptor distribution of Ribeye.

**Figure 7.**
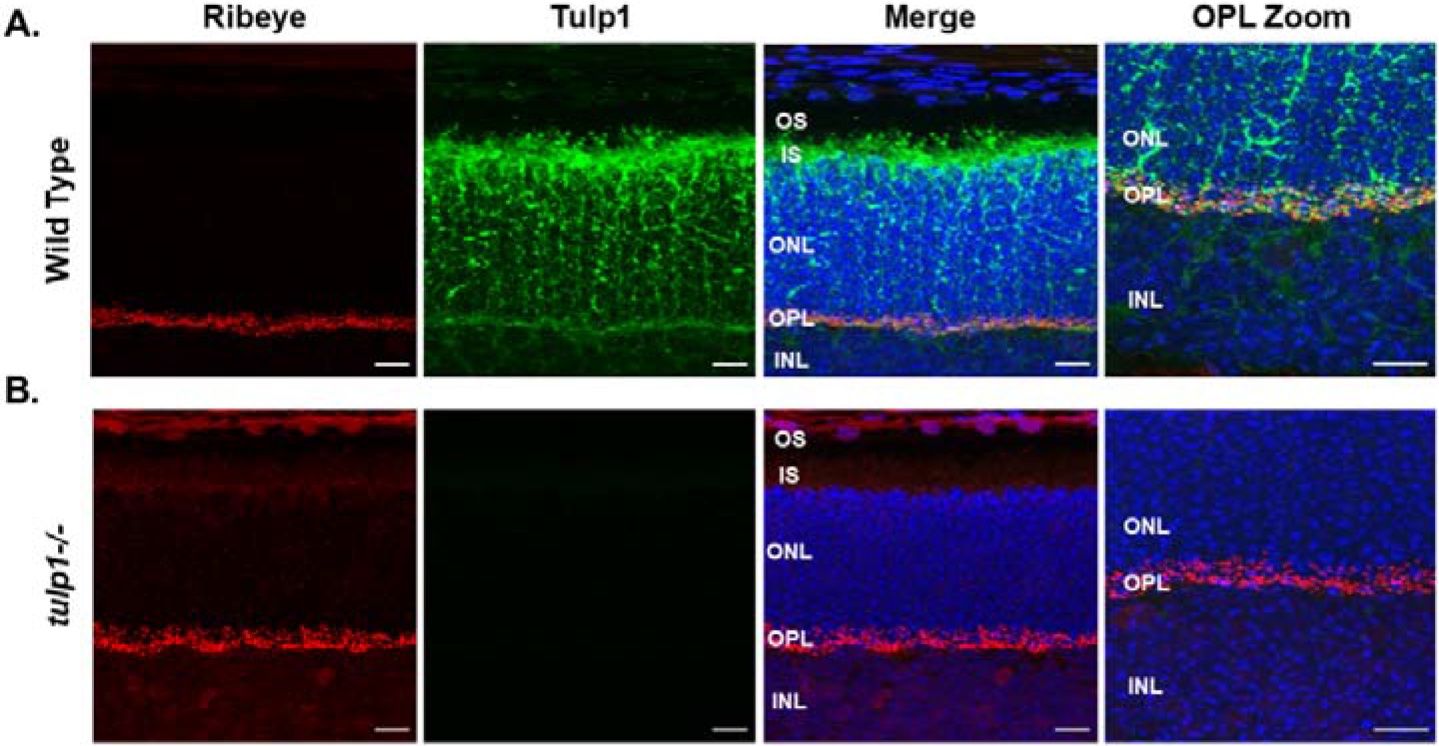
Immunolocalization of Ribeye and Tulp1 in P17 mouse retinas. (A) Wt retinal sections stained with Ribeye (red) and Tulp1 (green). (B) *Tulp1-/-* retinal sections stained with Ribeye (red) and Tulp1 (green). Sections were counterstained with DAPI (blue). Scale bar: 50 μm. INL, inner nuclear layer; OPL, outer plexiform layer; ONL, outer nuclear layer; IS, inner segment layer; OS, outer segment layer.

## 3. Discussion

Photoreceptor cells exhibit morphologically distinct compartments. At opposite ends of the cell, vesicular protein transport is required to coordinate the extreme bio-synthetic demands of phototransduction and neurotransmission. In the present study, we exploited the distinct photoreceptor compartments combined with a proteomic approach to demonstrate an interaction between Tulp1 and unique compartment-specific proteins. Our data indicate that Tulp1 has two distinct photoreceptor interactomes, one in the IS and one in the synapse.

We and others have previously identified Tulp1 interacting proteins in mice [34–37]. Tulp1 is a member of a family of related proteins named tubby-like proteins (Tulps) and encompasses the founding member Tub and the related Tulps, Tulp1 – 4. [38,39]. Mutations in human TULP1 have been associated with early-onset, severe retinitis pigmentosa (RP) and Leber congenital amaurosis (LCA) [40–44]. Patients harboring TULP1 mutations report early-onset night blindness, and severe visual acuity loss involving both rod and cone photoreceptors. Each Tulp family member is characterized by a signature carboxy-terminal “tubby” domain. A small motif in this domain binds selective phosphoinositides (PIPs), linking Tulps to the plasma membrane [36,45]. The amino terminus of Tulp proteins is diverse and directs distinct functions. For example, a short 20 amino acid sequence in the Tulp3 amino terminus binds to IFT-A, a complex important in retrograde protein transport in cilia [46]. This motif is not conserved in Tulp1 and therefore, not surprisingly, no IFT-A proteins were identified in our study. Our previous work has shown that Tulp1 can also associate with specific cellular membrane PIPs and interact with the cytoskeleton protein Actin, and the neuronal-specific GTPase Dynamin-1, all of which are involved in vesicular transport [34,35].

Herein, we demonstrate that Tulp1 interacts with MAP1B in both the photoreceptor IS and synaptic layers. MAP1B is primarily expressed in neurons and is a scaffold protein supporting intracellular protein transport systems and maintaining cellular structural integrity through microtubule stabilization and regulation [23]. The architecture of the photoreceptor requires interactions between cytoskeletal elements such as actin and protein transport such as the synaptic ribbon for neurotransmission and the axoneme for OS-destined phototransduction proteins. Photoreceptors lacking Tulp1 show severe defects affecting protein trafficking at both ends of the cell followed by rapid degeneration [11–13]. Rhodopsin along with several other OS-bound phototransduction proteins and two Rab proteins involved in the delivery of rhodopsin-containing vesicles to the base of the CC are mistrafficked in *tulp1-/-* photoreceptors [15,16]. In the *tulp1-/-* synapse, the tight spatial relationship between the ribbon-associated proteins, Bassoon and Piccolo, are disrupted, and few intact ribbons are present [14]. The consequence of MAP1B loss in photoreceptors is less straight forward. Four different MAP1B knockout mice have been generated with various differences in their phenotypes. In most cases, homozygous mice die during embryogenesis or shortly following birth due to gross neuronal defects [23]. One study examined the retinal morphology in 4-month old heterozygous mice and revealed photoreceptor degeneration including OPL defects [47,48]. To directly determine the effects of MAP1B loss on photoreceptor protein trafficking, a cell-specific knockout will need to be generated. Our study indicates that Tulp1 is a cytoskeletal interactor in both the photoreceptor IS and synapse through binding MAP1B and is required for maintenance of the structural scaffolds essential for intracellular trafficking of proteins. Our results are strengthened by the co-localization of Tulp1 and MAP1B in wt photoreceptors and our previous data showing that Tulp1 binds and co-localizes with Actin, also a MAP1B binding partner [34,49].

We identified Kif3a as an IS-specific Tulp1 interacting protein. This interaction is further supported by reciprocal IP analysis and co-localization experiments in retinal sections. Kif3a is one of two motor subunits of the heterotrimeric Kinesin-2 microtubule-dependent motor complex that facilitates IFT transport of axonemal-building tubulin subunits and ciliary membrane protein trafficking [29,50]. In photoreceptors, Kinesin-2 transports cargo proteins from the IS through the CC to maintain the axoneme and OS components [51]. Although commonly accepted that Kinesin-2 transports IFT complexes and associated cargo anterograde, the idea that Kinesin-2 mediates rhodopsin transport is controversial. In mice, photoreceptor-specific knockout of Kif3a results in a retinal phenotype very similar to the *tulp1-/-* phenotype including rhodopsin and arrestin mislocalization and progressive photoreceptor cell death, suggesting that Kinesin-2/IFT complexes mediate opsin transport [11,12,52–54]. Conversely, a separate conditional Kif3a knockout in mouse rod photoreceptors surprisingly showed normal trafficking of rhodopsin as well as other OS membrane proteins even during rapid photoreceptor degeneration [55]. Furthermore, a retina-specific tamoxifen inducible deletion of Kif3a in adult mice showed normal rhodopsin localization before degeneration while the same deletion in embryonic mice resulted in lack of axoneme/CC formation [31]. It remains to be determined whether transition zone and centriolar proteins are disrupted in *tulp1-/-* photoreceptors. Interestingly, a subset of OS resident proteins retain their proper localization in *tulp1-/-* retinas as well as a small portion of rhodopsin [12,15]. It is possible that some, but not all, cililary proteins may traffic via IFT driven by Kinesin-2. This pathway may not be absolutely essential for all protein transport but may instead regulate their ciliary levels, as previously proposed [50]. Evidence from numerous studies suggest that the mechanisms underlying protein trafficking in and out of cilia are complex and likely involves multiple processes, including diffusion, localization signals, and IFT motor-driven pathways [56,57].

Kif3a has also been reported to be present at the photoreceptor ribbon synapse [58]. However, we did not detect Kif3a in our synaptic-specific Tulp1 interactome. There are several possible explanations for our findings. First, the detection of protein protein interactions can be affected by various factors that may not be present in the OPL. Kif3a may have a weak binding affinity or a transient interaction with Tulp1, or either protein may require post-translational modifications in order to interact. Second, the other components of the Kinesin-2 motor complex are not expressed in photoreceptor terminals questioning the functional significance of Kif3a at the synapse [28,59]. In fact, there is evidence that presynaptic transport vesicles may use different Kinesin family members [60]. And lastly, evidence suggests that many presynaptic cargos share common vesicles as opposed to each protein using a unique set of vesicles [61,62]. For example, synaptic vesicle proteins are incorporated into synaptic vesicle protein transport vesicles while cytomatrix active zone proteins are associated with piccolo-bassoon transport vesicles.

Ribeye was identified as a Tulp1 interactor in our proteomic analysis of the photoreceptor synaptic-specific compartment. This is not surprising given our previous work demonstrating that Tulp1 is expressed in the OPL of photoreceptors and lack of Tulp1 causes abnormal ribbon architecture as well as bipolar cell dendritic malformation [14]. A separate study has shown that Tulp1 interacts with Ribeye via yeast two-hybrid experiments and mediates localization of the endocytic machinery at the periactive zone of photoreceptor synapses [37]. An endocytic role for Tulp1 is supported by our previous results identifying an interaction between Tulp1 and Dynamin-1, the mechanoenzyme that mediates periactive zone endocytosis [35]. In the current study, our photoreceptor synaptic-specific proteomic analysis confirms a direct Tulp1-Ribeye interaction in their native cellular compartment and IHC experiments show that Tulp1 and Ribeye co-localize to the OPL. As expected, Ribeye knockout abolishes all photoreceptor synaptic ribbons and causes a loss of synaptic vesicles in the vicinity of the active zone of photoreceptor ribbon synapses [32]. However, phototransduction in the OSs is unaffected [63]. Whether Tulp1, Dynamin-1 and other Ribeye interacting proteins are present and correctly localized in Ribeye-/- photoreceptors has yet to be determined.

From our findings, Tulp1 could function in several capacities, none of which are mutually exclusive. First, it may serve as an adapter protein involved in selecting cargo for inclusion into transport vesicles. This has been demonstrated for Tulp3, which has been functionally linked to phosphoinositide 4,5-bisphosphate (PI(4,5)P2) and the IFT-A complex through specific binding motifs to traffic membrane proteins to neuronal cilia [46,64]. Tulp1 may play a similar role in photoreceptors, as we’ve previously proposed [15,16]. The molecular function of Tulp1 as an adapter protein likely depends on its bi-partite structural organization bridging interactions between different proteins. Second, it may be part of a dynamic microtubule scaffold connecting transport vesicles with the cytoskeleton. In support of this view, our previous and current data indicate that Tulp1 binds multiple cytoskeletal components including Actin, MAP1B and Kif3a. It is known that Kinesin microtubule interactions can be modulated through the action of MAPs to achieve compartmentalized cargo distribution [24,65,66]. Tulp1 may act as a bridge molecule in regulating the correct delivery of cargo protein through binding both the compartment-specific cytoskeletal scaffold proteins and motor proteins. A recent bioinformatics analysis also predicted several cytoskeletal Tulp1 interactors [67]. Lastly, Tulp1 may regulate vesicle trafficking from the trans-Golgi network to the base of the CC through the endosomal pathway, as has been proposed for rhodopsin carrier vesicles [68]. This possibility is less likely as this trafficking route has not been established for other OS-specific carrier vesicles or synaptic-destined vesicles.

Understanding the molecular processes that facilitate protein movement is vital to photoreceptor cell biology considering the enormous amount of proteins that are synthesized and delivered to the OS and the continuous vesicular cycling of neurotransmitter release that occurs at the synaptic terminal. Our findings suggest that Tulp1 is required in multiple photoreceptor compartments and interacts with compartment-specific and compartment-non-specific proteins. This complexity may explain the severe phenotype of TULP1-induced retinal degeneration. It is only through a complete understanding of the photoreceptor compartment-specific function of Tulp1 that therapeutic strategies can be developed to treat this severe retinopathy. The challenge will clearly become how to treat a photoreceptor degeneration with multiple sites of dysfunction. Whether the described interactions are lost when Tulp1 is mutated as compared to knocked out is unknown; however, we are currently generating several knock-in mouse models harboring RP-associated *TULP1* mutations to investigate these questions. In summary, our proteomic analysis of Tulp1 interactors suggests photoreceptor compartment-specific utility of this multifaceted protein.

## 4. Materials and Methods

### 4.1 Animals

The generation of *tulp1-/-* mice has been described previously [11]. Wild-type (wt) C57Bl6/J mice were purchased from the Jackson Laboratory and Long Evans rats were purchased from Charles River Laboratory. All animals were housed in the Cole Eye Institute Biological Resources Unit at the Cleveland Clinic. All animal experiments were approved by the Institutional Animal Care and Use Committee of the Cleveland Clinic and were performed in compliance with the ARVO Statement for the Use of Animals in Ophthalmic and Visual Research.

### 4.2 Serial Tangential Sectioning

Tangential sectioning of rat retinas were carried out as previously described with several optimizations [69]. Retinas from adult Long Evans rats were dissected in DMEM/F12 media supplemented with complete protease inhibitors (Roche, Indianapolis, IN) and positioned above a disc of nitrocellulose disc oriented photoreceptor-side down. Each retina was cut into halves or quarters and flattened individually in a custom-made chamber by positioning the retina and nitrocellulose disc above a glass capillary array (BURLE Electro-Optics, Sturbridge, MA) and slowly removing the media from the lower chamber using a hand-drawn 10 ml syringe attached to the bottom chamber with a Luer lock fitting. The flattened retina was secured onto a 2×2 cm^2^ glass slide using superglue and covered with a slide wrapped in non-stick optically-clear tape, separated from the bottom slide by 0.5 mm plastic spacers. This flat mount preparation was clamped on each slide with small binder clips and placed on dry ice to freeze for ~30 min. A mound of OCT compound (Sakura Finetek USA, Torrance, CA) was frozen on a cryostat chuck and sectioned to create a flat surface large enough to accommodate the bottom glass slide. After removing the clamps and top slide, the bottom slide was held against the OCT surface and frozen in place by the addition of water drops around the back of the slide base. The peripheral edges of each retina were trimmed using a scalpel blade to remove any folded or uneven areas. Finally, the retina was serial sectioned at a thickness of 10 μm. Each section was collected in 50 μl of Tris-Glycine SDS sample buffer (Life Technologies, Carlsbad, CA) or Pierce IP lysis buffer (Thermo Scientific, Rockford, IL) depending upon downstream analysis. Each serial sectioned retina yielded between 20 and 30 sections, and in total are considered an experimental “set”. Sample sets were stored at −80°C until analyzed.

### 4.3 Optical Coherance Tomography (OCT)

To assess the quality of the dissected retina, OCT images of retinal flat mount preparations were collected using a Model 840HR SDOIS (Bioptigen, Inc, Morrisville, NC) with a center operating wavelength of ~840 nm and axial in-depth resolution of 6-7 μm. Prior to freezing, retina flat mount preparations were placed on a rotational mount and angled at ~7 degrees with respect to the central imaging axis as to avoid signal saturation from air/glass/tissue interface reflections. Acquired OCT images of the preparation included both single B-scans (1000 A-scans/B-scan) and entire 3-D volumes (500 B-scans/volume x 100 A-scans/B-scan). Averaging (line and frame) was employed to enhance image contrast and minimize background noise.

### 4.4 Dot-blot Analysis

To assess the quality of tangential sectioning, 0.5 μl aliquots of each tangential section in a set was dotted onto a dry nitrocellulose membrane, blocked with Odyssey Blocking Buffer (LI-COR Biosciences, Lincoln, NE) and probed with antibodies against rhodopsin (1D4 1:5000, Thermo Scientific). After incubation with IRDye 800CW goat anti-mouse secondary antibody (LI-COR), the dot blot membrane was imaged on an Odyssey infrared scanner (LI-COR). Only the tangentially sectioned retinas which stain positive for rhodopsin in the outermost three to six sections by dot-blot, corresponding to the photoreceptor OS, were selected for further analysis.

### 4.5 SDS-PAGE and Western Blot Analysis

Equal volumes of tangentially sectioned sets were reduced, denatured, and separated on 4-20% Tris-Glycine SDS-PAGE gels (Life Technologies) and transferred overnight onto polyvinylidene fluoride (PVDF) membranes. Membranes were blocked with 5% blotting grade blocker (Bio-Rad, Hercules, CA), incubated with primary antibody at 4°C overnight, washed, and then incubated with peroxidase-conjugated secondary antibody. Chemiluminescent detection was achieved using Western Lightning ECL Plus or Pro (PerkinElmer, Waltham, MA) followed by film exposures. In some experiments, membranes were stripped using Restore Western Blot stripping buffer (Thermo Scientific), re-blocked, and re-probed with a different primary antibody. Retinal layer-specific primary antibodies were: Tulp1 (M-tulp1N; 1:2000)[11], ubiquitous mitochondrial creatine kinase (uMtCK C-18, Santa Cruz, 1:500), synaptophysin (SVP38, Santa Cruz, 1:1000), phosducin (kindly provided by Dr. Sokolov, West Virginia University, 1:1000) [18], and rhodopsin (1D4, Thermo Scientific, 1:5000).

### 4.6 Immunoprecipitation (IP)

IS, OPL, and ganglion cell layer (GCL) compartments were identified by Western blotting using a reference set of tangentially sectioned retina. In order to obtain sufficient protein for downstream proteomic analysis, samples containing IS, OPL and GCL across multiple sets were pooled. As negative controls the GCL, an inner retinal layer which lacks Tulp1, and whole retina lysate from P17 *tulp1-/-* mice were also co-IPed. Each pooled region contained between 200-960 μg of total protein. Co-IP was performed using the Dynabeads-Protein A or Protein G Immunoprecipitation kit (Life Technologies, rabbit antibody with Protein A, mouse antibody with Protein G). Each pooled sample was precleared with non-specific rabbit IgG-bound magnetic beads for 1 hr at 4°C. For each co-IP experiment, magnetic beads were bound with ~10 μg rabbit polyclonal anti-Tulp1 antibodies (mTulp1N) and beads were cross-linked with 5 mM BS3 (Bis[sulfosuccinimidyl] suberate, Thermo Scientific) in 20 mM HEPES for 30 min. Following quenching of the cross-linking reaction and washing, compartment-specific lysates were added to the antibody-linked beads for 2 hours with rotation at 4°C. Tulp1 and interacting proteins were eluted in 20-40 μl Tris-Glycine SDS sample buffer. Reciprocal co-IPs were performed on whole rat retina lysate to confirm identified Tulp1-binding partners using the following target antibodies: MAP1B (NB100-68256, Novus Biologicals), Kif3a (EPR5087, Abcam), and Ribeye (612044, BD Transduction).

### 4.7 Liquid Chromatography Tandem Mass Spectrometry (LC-MS/MS)

Eluted IP products were separated on SDS-PAGE gels. For in-gel digestions, the lanes were excised and divided into a number of smaller areas for trypsin digestion according to a previously published method [70]. Trypsinized peptides were extracted from the polyacrylamide and resuspended in 1% acetic acid for analysis by LC-MS/MS on a LTQ-Obitrap Elite hybrid mass spectrometer system coupled to a Dionex Ultimate 3000 HPLC (Thermo Scientific). Five μL aliquots of the digests were loaded onto a 75 μm Acclaim Pepmap C18 reverse phase column (Thermo Scientific) and eluted by an acetonitrile/0.1% formic acid gradient. The digest was analyzed using a data dependent acquisition and the proteins were identified by searching the LC-MS/MS data with the programs Mascot and Sequest against the rat or mouse Reference Sequence Databases. These search results were uploaded into the program Scaffold (Proteome Software, Portland, OR) for relative quantitation using normalized spectral counts for each identified protein. For proteomic analysis, the relative quantity of identified proteins was determined by comparing the number of spectra, termed spectral counts, used to identify each protein. The numerical values used in quantitation corresponded to the normalized spectral counts reflecting the number of exclusive spectra/total spectral counts identified in the LC-MS/MS experiment. Identified interacting proteins were analyzed through the use of QIAGEN’s Ingenuity® Pathway Analysis (IPA®, QIAGEN, Redwood City, CA) and UniProt online protein knowledgebase (http://www.uniprot.org/).

### 4.8 Immunohistochemistry (IHC)

Mouse eye from P17 wt and *tulp1-/-* mice were enucleated and immediately frozen in OCT with liquid nitrogen and store at −80oC for sectioning the next day. Tissue was sectioned on an adhesive tape at 10-μm thickness using a CM1950 (w/Cryolane) cryostat (Leica, Wetzlar, Germany) at −30°C. Retinal sections were fixed in 4% paraformaldehyde (PFA) for 5 mins, followed by brief washing in 1X PBS. Tissue sections were permeabalized with 1X PBS containing 0.025% Triton X-100 (PBST) for 10 mins and then washed 3 times with 1X PBS, 5 mins each. Slides were then incubated with blocking solution (1% BSA and 10% Donkey serum in freshly prepared 1X PBS) for 2 hrs at RT and then subsequently incubated with primary antibodies: MAP1B at 1:100 (N-19, Santa Cruz Biotechnology), Kif3a at 1:300 (Abcam, ab11259), Ribeye at 1:1000 (BD Transduction, 612044) and Tulp1 at 1:500 (mTulp1) [11] (diluted in 1X PBS with 1% BSA) overnight at 4oC. Next day primary antibodies were removed and the slides were washed 3 times with PBST on a rocker for 5 mins each. Secondary antibodies used for detection were as follows: Alexa Fluor donkey anti-rabbit 594 and donkey anti-goat 488 (1:1000, Life Technologies) diluted in 1X PBS with 1% BSA and incubated in the dark for 1 hr at RT. Slides were then washed 3 times with PBST 10 mins each. One drop of mounting media containing DAPI (Vectasheild) was placed on the samples and mounted with coverslip. Slides were stored in the dark overnight and imaged using a fluorescence microscope (Zeiss Axio Imager. Z2, Germany).

## Author Contributions

Conceptualization, L.A.E. and S.A.H.; methodology, L.A.E., S.S., G.J.T.P., S.A.H.; writing—original draft preparation, L.A.E. and S.A.H.; writing—review and editing, S.S. and S.A.H.; funding acquisition, S.A.H. All authors have read and agreed to the published version of the manuscript.

## Institutional Review Board Statement

The study was conducted according to the guidelines of the Declaration of Helsinki, and approved by the Institutional Animal Care and Use Committee of the Cleveland Clinic, Protocol: 2019-2242; Date of approval: 9/27/2019.

## Informed Consent Statement

Not applicable.

## Data Availability Statement

In this section, please provide details regarding where data supporting reported results can be found, including links to publicly archived datasets analyzed or generated during the study. Please refer to suggested Data Availability Statements in section “MDPI Research Data Policies” at https://www.mdpi.com/ethics. You might choose to exclude this statement if the study did not report any data.

## Acknowledgments

We thank Dr. Neal S. Peachey for insightful comments on the manuscript.

